# LoReMINE: Long Read-based Microbial genome mining pipeline

**DOI:** 10.64898/2026.02.02.703231

**Authors:** Amay A. Agrawal, Chantal D. Bader, Ronald Garcia, Rolf Müller, Olga V. Kalinina

**Affiliations:** Department of Drug Bioinformatics, Helmholtz Institute for Pharmaceutical Saarland (HIPS) / Helmholtz Centre for Infection Research (HZI), Saarbrücken, Saarland, Germany; Graduate School of Computer Science, Saarland University, Saarbrücken, Saarland, Germany; Department of Microbial Natural Products, Helmholtz Institute for Pharmaceutical Saarland (HIPS) / Helmholtz Centre for Infection Research (HZI), Saarbrücken, Saarland, Germany; PharmaScienceHub, Saarbrücken, Saarland, Germany; German Center for Infection Research (DZIF), Partner Site Hannover-Braunschweig, 38124 Braunschweig, Germany; Faculty of Medicine, Saarland University, Homburg, Saarland, Germany; Center for Bioinformatics, Saarland University, Saarbrücken, Saarland, Germany

## Abstract

Microbial natural products represent a chemically diverse repertoire of small molecules with major pharmaceutical potential. Despite the increasing availability of microbial genome sequences, large-scale natural product discovery remains challenging because the existing genome mining approaches lack integrated workflows for rapid dereplication of known compounds and prioritization of novel candidates, forcing researchers to rely on multiple tools that requires extensive manual curation and expert intervention at each step. To address these limitations, we introduce LoReMINE (**Lo**ng **Re**ad-based Microbial genome **min**ing pipelin**e**), a fully automated end-to-end pipeline that generates high-quality assemblies, performs taxonomic classification, predicts biosynthetic gene clusters (BGCs) responsible for biosynthesis of natural products, and clusters them into gene cluster families (GCFs) directly from long-read sequencing data. By integrating state-of-the-art tools into a seamless pipeline, LoReMINE enables scalable, reproducible, and comprehensive genome mining across diverse microbial taxa. The pipeline is openly available at https://github.com/kalininalab/LoReMINE and can be installed via Conda (https://anaconda.org/kalininalab/loremine), facilitating broad adoption by the natural product research community.

**Author summary:** For decades, microbial natural products have been a major source of medicines, with most of the clinically used antibiotics being their derivatives. Recent advances in DNA sequencing technologies now allow the reconstruction of more complete and continuous microbial genomes, revealing a vast and largely untapped diversity of biosynthetic gene clusters responsible for natural product biosynthesis. Despite these advances, large-scale natural product discovery remains difficult because current genome mining approaches rely on many separate tools and lack an integrated workflow to dereplicate known compounds and prioritize novel biosynthetic pathways. To address these limitations, we introduce LoReMINE, an automated pipeline designed to simplify microbial genome mining directly from long-read sequencing data. LoReMINE integrates genome assembly, taxonomic classification, identification of biosynthetic gene clusters, and their clustering into gene cluster families within a single, reproducible workflow. This streamlined approach enables scalable analysis across diverse microbial taxa and facilitates comprehensive exploration of microbial biosynthetic potential. The pipeline is designed for both experimental and computational researchers, helping to advance natural product research and contribute towards the discovery of new therapeutic drugs.

## Introduction

Natural products (NPs) have historically been an important source for new therapeutic drugs. They are small molecules produced by plants, microbes, and fungi that are not directly essential for growth and development, but they help the producing organism in adapting to its environment or serve as a defense mechanism, thereby supporting its survival [1]. One of the most famous example of a natural product is the antibiotic penicillin, which was derived from the fungus *Penicillium notatum* [2]. Some of the other widely prescribed drugs that are either natural products or their derivatives include aspirin (anti-inflammatory), doxorubicin (anti-cancer), amphotericin B (antifungal), oseltamivir (antiviral), acarbose (antidiabetic), avermectin (antiparasitic), tetracycline (antibiotic), etc. [3, 4]

Traditional approaches for discovering new NPs primarily relied on the biological screening of crude extracts to isolate active compounds [5]. While these methods proved effective in identifying numerous bioactive NPs, they were constrained by certain limitations. Many NPs are poorly expressed under standard laboratory conditions, making them undetectable through conventional techniques, and as a result, a vast pool of potentially valuable compounds remains inaccessible. Additionally, these approaches are labor-intensive, have low-throughput and are often hindered by the frequent rediscovery of already known molecules, thereby reducing their efficiency in uncovering novel bioactive substances [6].

As the cost of whole-genome sequencing has continuously declined over the last few decades, the focus has now shifted from traditional bioactivity-guided approaches towards genome mining based approaches for finding new NPs [7, 8]. Genome mining based approaches are based on the fact that NPs are typically synthesized by groups of enzymes, whose genes are encoded in close proximity in the genome, forming a so-called biosynthetic gene clusters (BGCs) that can be effectively found computationally. These BGCs are categorized into distinct classes based on their core enzymes and assembly logic, with major classes including non-ribosomal peptide synthetases (NRPSs), polyketide synthases (PKSs), ribosomally synthesized and post-translationally modified peptides (RiPPs), terpenes, and saccharides [9].

A key advantage of genome mining is its ability to detect “cryptic” or “silent” BGCs that are poorly expressed under standard laboratory conditions and therefore remain inaccessible via traditional methods. Once identified, these hidden pathways can be activated using strategies such as heterologous expression, unlocking a large reservoir of previously hidden chemical diversity [10]. Genome mining also allows targeted drug discovery by searching for specific genomic markers such as self-resistance genes or bioactive chemical features to identify natural products with desired biological activities [8]. Furthermore, combining genome mining with taxonomic classification provides an evolutionary framework for prioritizing BGCs from rare or underexplored microbial lineages, which are more likely to yield novel compounds and also help to reduce the rediscovery of known natural products [11, 12].

Genome mining based NP discovery involves multiple steps, including genome assembly, identification of BGCs, comparative genomic analysis to identify similarities and differences between BGCs and organisms, and prediction of biosynthesized natural product scaffolds [13]. Despite its transformative potential, genome mining is strongly dependent on the quality of genome assemblies. Short-read sequencing technologies, such as Illumina, has been largely responsible for the rapid accumulation of draft genome assemblies at the National Centre for Biotechnology Information (NCBI), but they often yield highly fragmented assemblies. As a result, BGCs encoding large multimodular enzymes such as PKSs and NRPSs are often splitted across multiple regions, making it difficult to detect and interpret them correctly. To alleviate this, long-read sequencing platforms, such as PacBio and Oxford Nanopore Technologies (ONT), have become indispensable in genome mining studies, as they enable the generation of contiguous assemblies that span complete BGCs, providing an unprecedented opportunity to fully exploit microbial genomes for NP discovery.

Once a high-quality assembly is obtained, the next step is to identify the BGCs. Several tools like antiSMASH [14], PRISM [15], DeepBGC [16] have been developed to identify BGCs, among which antiSMASH is regarded as a gold standard tool for this purpose. It is a rule-based tool which uses profile hidden Markov models (pHMMs) to identify BGCs, and currently it contains detection rules for identifying 101 different classes and sub-classes of BGCs.

To enable exploration beyond single-genome studies, antiSMASH database [17] provides a large-scale repository of precomputed BGCs across thousands of microbial genomes, offering a valuable resource for comparative analyses across different taxa. Further, many of these predicted BGCs have also been experimentally linked to specific natural products with characterized chemical structures, giving rise to a database called Minimum Information about a Biosynthetic Gene cluster (MIBiG [18]) that currently contains more than 2500 validated gene cluster–molecule pairs. Given the enormous diversity of BGCs across these databases, comparison tools, such as BiG-SCAPE [19] and BiG-SLiCE [20], have been developed. They facilitate mapping relationships between thousands of BGCs by clustering them into gene cluster families (GCFs). BGCs belonging to different GCFs are expected to produce chemically different compounds, while the products of BGCs from the same GCF can be quite similar. The clustering output readily allows the user to identify GCFs with known members from MIBiG, while, on the other hand, GCFs with no MIBiG reference BGCs may point to biosynthetic pathways that are completely novel and unexplored.

Despite the availability of these powerful tools and databases for genome mining, a unified workflow that integrates them for large-scale natural product discovery is still missing. To address this critical gap, we present LoReMINE (**Lo**ng **Re**ad-based Microbial genome **min**ing pipelin**e**), an end-to-end workflow for microbial natural product discovery directly from long-read sequencing data. LoReMINE integrates multiple modules into a unified pipeline: *(i) de novo* genome assembly using multiple long-read assemblers with automated selection of the highest quality assembly, (ii) taxonomic classification, (iii) BGC detection, and (iv) clustering of identified BGCs into GCFs to facilitate comparative analysis across different databases and taxa. By combining these steps into a unified workflow, LoReMINE facilitates comprehensive exploration of natural product diversity which has the potential to yield novel drug candidates.

## Materials and methods

LoReMINE is designed as a fully automated, modular workflow to enable end-to-end discovery of microbial natural products directly from long-read sequencing data. The workflow integrates four core submodules: (i) assemble, (ii) taxonomy, (iii) identify_bgcs, and (iv) bgc_clustering, each of which can be executed independently or as part of the complete workflow (Figure 1). The individual submodules are described below.

**Fig 1.**
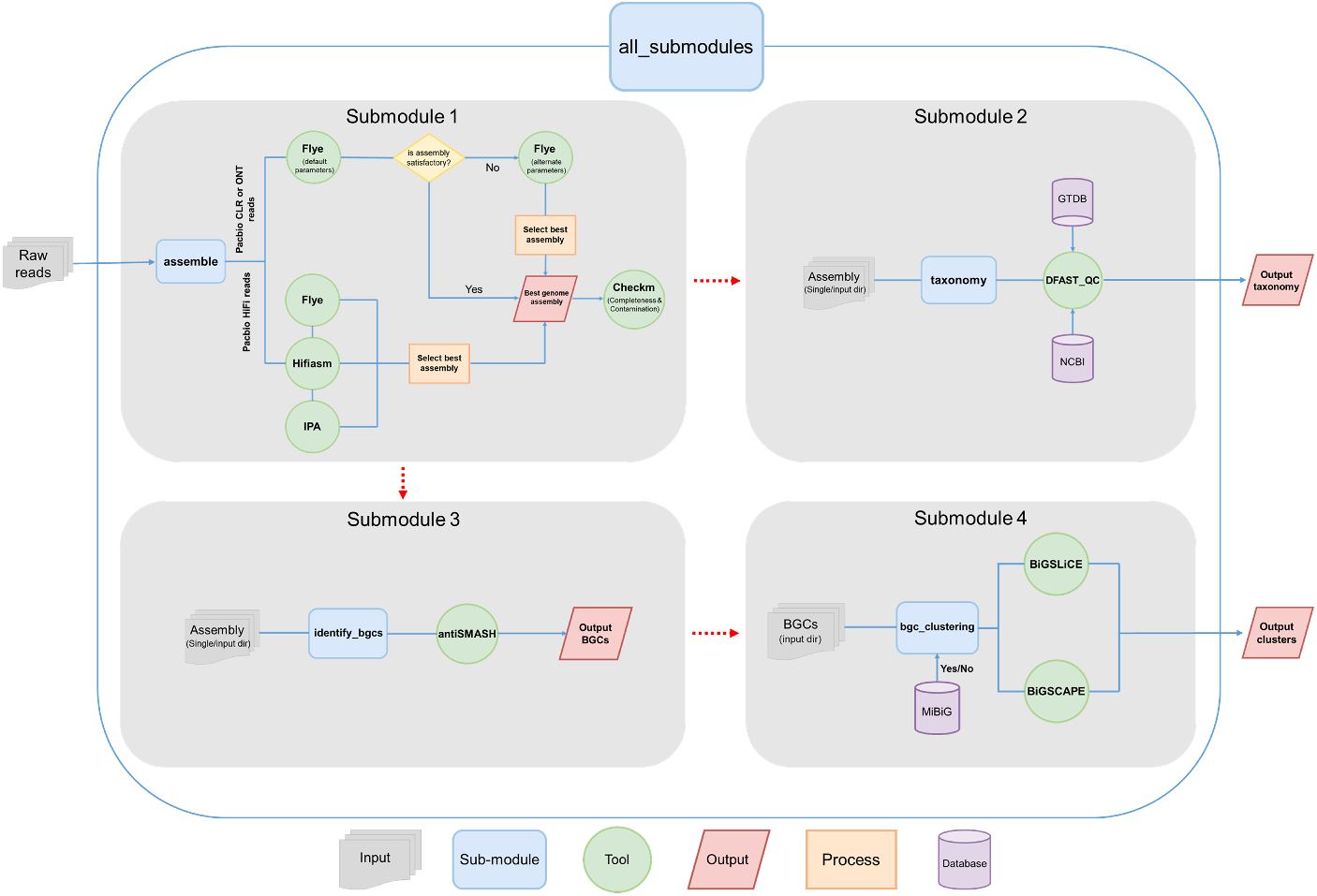
Workflow of the LoReMINE pipeline. LoReMINE consists of four submodules that can be run individually or as a complete workflow. Submodule 1 takes sequencing reads as input and, depending on the read type, applies an appropriate assembly strategy to generate the best-quality genome assembly. The resulting assembly can then be used as input for Submodule 2, which performs taxonomic classification of the input genome, and Submodule 3, which identifies BGCs in the input genome. The predicted BGCs, optionally combined with reference BGCs from the MIBiG database, are provided to Submodule 4, which clusters BGCs into GCFs. The resulting GCFs can be visualized to distinguish known biosynthetic pathways from potentially novel ones. Alternatively, the `“all_submodules”` option provides a fully automated workflow that takes sequencing reads as input, executes all four submodules sequentially, and directly produces GCF outputs without requiring any manual intervention.

### assemble

This submodule performs *de novo* genome assembly from long-read sequencing data generated by PacBio (both HiFi and continuous long reads (CLR)) and Oxford Nanopore Technologies (ONT) platforms. The pipeline supports both single-strain assembly and batch processing of multiple strains in a single run, enabling scalable analysis of large sequencing datasets. To maximize assembly quality and minimize assembler bias, LoReMINE integrates multiple state-of-the-art long-read assemblers.

For PacBio HiFi reads, three variants of assemblies are independently generated using Flye [21], Hifiasm [22], and IPA [23]. For PacBio CLR and ONT reads, the assembly is first performed using Flye with default parameters. If the resulting assembly is deemed unsatisfactory by the user, the parameter (`--alt_param`) can be activated, which performs additional assemblies by adjusting the “minimum overlap” parameter of the Flye assembler. Each candidate assembly undergoes a quality control step performed using DustMasker [24, 25] to remove contigs having low complexity regions such as homopolymer runs and simple sequence repeats (SSRs).

To automatically select the best genome assembly among all candidates, the pipeline employs a weighted scoring algorithm that integrates multiple quality metrics, including chromosome circularity, number of circular contigs, number of contigs, and contiguity (N50). Each assembly is evaluated across these metrics, and an assembly score is computed as described in Listing 1. The weights assigned to each metric are provided as default values but can be customized by the user as needed. Briefly, the procedure with default weights assign a very high score to assemblies containing a putative circular chromosome, defined as a contig representing at least 90% of the expected genome size (default: 5 Mbp). Additional circular contigs contribute minor positive scores, whereas fragmented assemblies incur strong penalties in the overall score. Continuity is also rewarded, with higher N50 values receiving a proportional bonus.

Following the scoring step, the pipeline generates a summary file `“chosen_best_assembly.txt”`, which lists all assemblies along with their assembly statistics and ranking from best to worst (based on their computed score). The top-ranked assembly with the highest overall weighted score is automatically selected as the final genome assembly for downstream analyses. In addition, the completeness and contamination of the top-ranked assembly are also evaluated using Checkm [26] and included in this file to assist users in assessing the final assembly quality. However, users can manually review and easily identify alternative assemblies using the detailed summary information provided in this file **(Supplementary Figure 1 and 2)**. This multi-assembler, score-based approach ensures that LoReMINE consistently produces high-quality, structurally complete genome assemblies across diverse long-read datasets.

**Listing 1.**
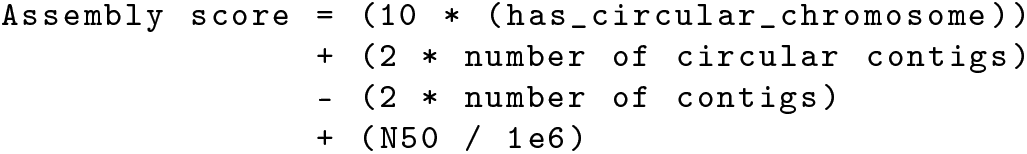
Weighted scoring formula used for assembly selection.

### taxonomy

This submodule performs automated taxonomic classification of input genomes using the DFAST_QC tool [27]. It supports two database configurations: a compact version (<1.5 GB) optimized for fast classification on resource-limited systems, and a full version (>100 GB) that provides a comprehensive coverage across bacterial taxa. Users can select either database configuration during setup, depending on their available resources and desired classification depth. DFAST_QC utilizes both the NCBI Reference Database [28] and Genome Taxonomy Database [29] reference taxonomies to ensure consistent, reliable, and up-to-date assignments.

Upon completion of taxonomic analysis, the pipeline automatically integrates the outputs from both databases and consolidates them into a single user-friendly summary file called `“identified_taxonomy.txt”` **(Supplementary Figure 3, 4, 5, and 6)**, providing a unified view of the genome’s taxonomic identity and facilitating downstream comparative or diversity analyses.

### identify_bgcs

This submodule detects BGCs in input genomes using the antiSMASH tool [14]. antiSMASH identifies a wide range of BGC types including polyketide synthases (PKS), nonribosomal peptide synthetases (NRPS), terpenes and others, and currently incorporates detection rules for 101 distinct BGC types. The output of this submodule consists of all the BGCs predicted by antiSMASH, along with their genomic locations, BGC class assignments, and detailed annotation reports, all of which can be interactively explored and visualized in a web browser and used as an input for subsequent analyses within the pipeline.

### bgc_clustering

This submodule is used for clustering the input BGCs to identify the gene cluster families (GCFs) that facilitate comparative and evolutionary analyses across both small and large-scale genomic projects. Clustering can be performed with both BiG-SLiCE (default threshold = 0.4) [20] and BiG-SCAPE (default threshold = 0.5) [19] using either the default or a user-defined threshold. The submodule also allows users to include reference BGCs from the MIBiG database [18], enabling a direct comparison between predicted and experimentally described BGCs. This integration helps to identify novel or unique GCFs that may represent previously uncharacterized natural product biosynthetic pathways.

Following clustering, the pipeline generates a comprehensive summary file called “output_clusters.tsv” for both tools, which lists all BGCs along with their assigned GCFs, similarity scores, BGC completeness, and length of the BGCs. Additionally, BiG-SCAPE output is provided as an interactive HTML visualization that enables users to explore cluster networks, inspect relationships among GCFs, and intuitively assess novelty and diversity within their dataset.

### all_submodules

This submodule enables execution of the entire pipeline in a single run. It supports both single-strain and batch-mode execution, allowing users to analyze individual isolates or large collections of strains simultaneously. In the batch mode, the pipeline automatically identifies the best genome assemblies for all strains, determines their taxonomy, detects BGCs in each best identified genome assembly, and subsequently performs clustering of identified BGCs across all strains to delineate GCFs.

Additionally, users may include MIBiG reference BGCs for the clustering step to facilitate comparative analysis and identify which GCFs correspond to known natural product families versus potentially novel clusters. Thus, this submodule provides a fully automated, end-to-end solution for high-throughput microbial natural product genome mining directly from the long-read sequencing data.

## Results

To illustrate how LoReMINE facilitates microbial natural product genome mining directly from long-read sequencing data, we applied the complete pipeline to ten bacterial strains, comprising six novel isolates and four type strains, generated using different long-read sequencing technologies. This use case highlights the pipeline’s end-to-end capabilities, automating assembly selection, taxonomic assignment, BGC detection, and GCF clustering. The full workflow was executed using the **all_submodules** option (Listing 2), and the outputs generated by each submodule are presented below.

**Listing 2.**
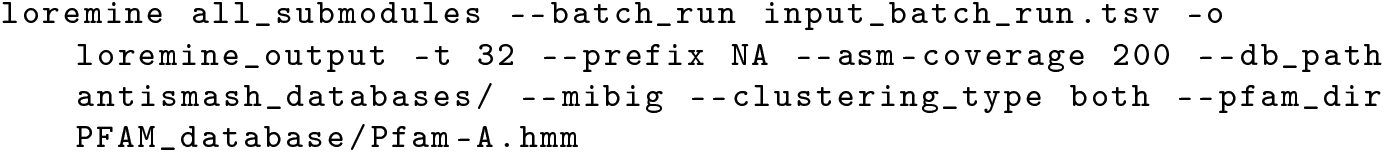
Running LoReMINE with all_submodules.

### assemble

The pipeline began by assembling ten strains sequenced using a mixture of PacBio CLR, PacBio HiFi, and Oxford Nanopore Technologies (ONT) long-reads. Leveraging the assemble submodule, we were able to reconstruct high-quality genomes for all strains (Table 1). Nine of the ten genomes were assembled into single circular contigs, while one genome yielded two circular contigs, the smaller of which, only *∼* 2.5 kb, may represent a plasmid, mobile genetic element intermediates, phage-derived elements, or assembly artifacts. All assemblies contained circular chromosomes, reflecting complete or near-complete genome reconstruction. For four type strains (DK1622^T^, DSM43024^T^, DSM43181^T^, and DSM43747^T^), the pipeline produced assemblies that were either comparable to or improved upon previously published assemblies in the literature (NCBI accessions: ASM1268v1, ASM30849v1, ASM1464805v1, and SREG). Notably, the assemblies for which improvements were observed had previously been generated using short-read sequencing, whereas LoReMINE leveraged long-read data, resulting in increased contiguity and improved assembly quality. Further quality assessment showed that each assembly exhibited over 90% completeness and less than 10% contamination, confirming that the pipeline consistently produced high-quality assemblies across all strains.

**Table 1.**
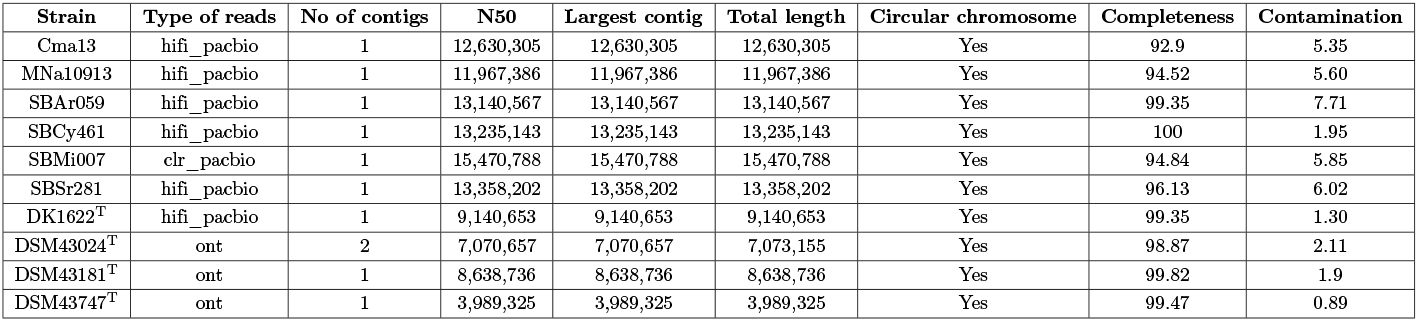
Genome assembly statistics for all ten bacterial strains. ^T^ denotes type strains.

### taxonomy

Following the reconstruction of high-quality genomes, the pipeline proceeded to determine the taxonomy of all ten strains. As described earlier, the taxonomy submodule integrates classification results from both the NCBI and the GTDB reference databases, enabling a comprehensive taxonomic assessment. Among the analyzed genomes, six strains represent novel isolates, and for these genomes, the average nucleotide identity (ANI) does not exceed the 95% threshold typically required for confident species-level classification. As a result, the pipeline was unable to definitively assign species-level taxonomy for each of these six strains (Table 2). Nevertheless, for each strain, at least one of the two databases successfully provided a nearest species-level classification, even when formal classification criteria were not met. In contrast, the remaining four type strains correspond to previously described organisms in the literature, and for them, ANI surpassed the 95% threshold. For these genomes, the taxonomic assignments produced by the pipeline were consistent with the species annotations reported in NCBI, demonstrating the accuracy of our pipeline.

**Table 2.**
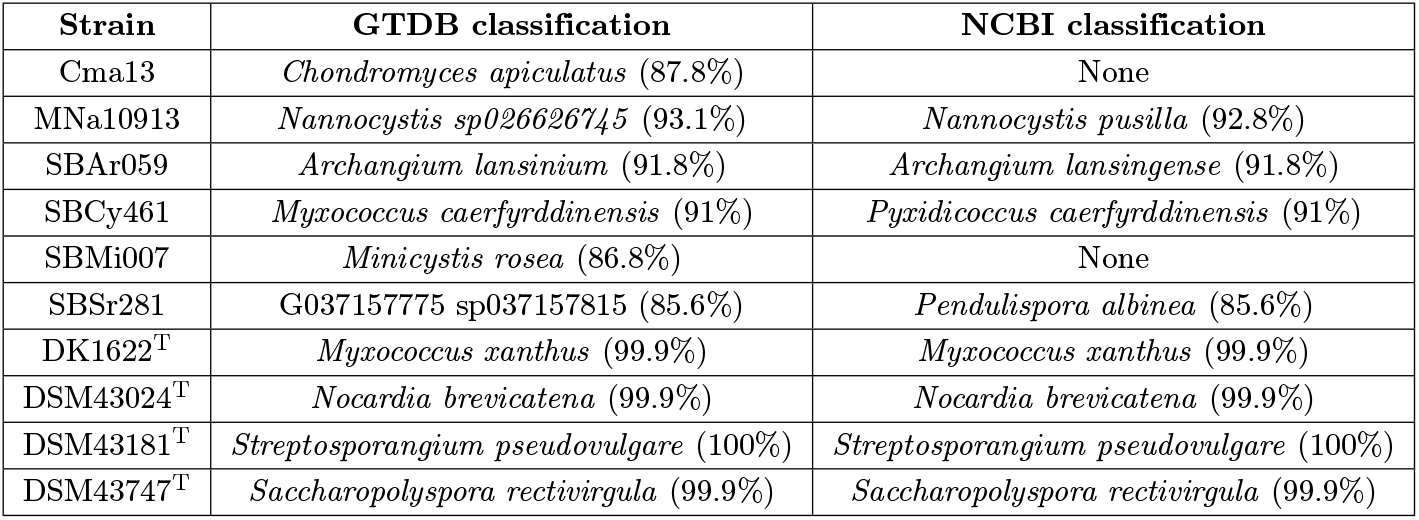
Species-level taxonomic classification of ten bacterial strains. Values shown in parentheses denote the ANI (in %) to the closest matching reference genome within each respective database. ^T^ denotes type strains.

| Strain | GTDB classification | NCBI classification |
| --- | --- | --- |
| Cma13 | *Chondromyces apiculatus* (87.8%) | None |
| MNa10913 | *Nannocystis sp026626745* (93.1%) | *Nannocystis pusilla* (92.8%) |
| SBAr059 | *Archangium lansinium* (91.8%) | *Archangium lansingense* (91.8%) |
| SBCy461 | *Myxococcus caerfyrddinensis* (91%) | *Pyxidicoccus caerfyrddinensis* (91%) |
| SBMi007 | *Minicystis rosea* (86.8%) | None |
| SBSr281 | G037157775 sp037157815 (85.6%) | *Pendulispora albinea* (85.6%) |
| DK1622<sup>T</sup> | *Myxococcus xanthus* (99.9%) | *Myxococcus xanthus* (99.9%) |
| DSM43024<sup>T</sup> | *Nocardia brevicatena* (99.9%) | *Nocardia brevicatena* (99.9%) |
| DSM43181<sup>T</sup> | *Streptosporangium pseudovulgare* (100%) | *Streptosporangium pseudovulgare* (100%) |
| DSM43747<sup>T</sup> | *Saccharopolyspora rectivirgula* (99.9%) | *Saccharopolyspora rectivirgula* (99.9%) |

### identify_bgcs

To assess the biosynthetic capacity of these genomes, the pipeline proceeded to identify the BGCs for all strains. In total, 389 BGCs were detected across all strains, and only two of these were fragmented. This underscores the importance of generating high-quality, near-complete assemblies, as fragmented genomes often yield incomplete BGCs, making it very challenging to reliably link metabolites to their corresponding clusters and thereby limiting genome-guided drug discovery. SBAr059 and DSM43747^T^ contain the highest and lowest number of BGCs (61 and 10), respectively (Figure 2A). Although non-ribosomal peptides (NRPs) and polyketides (PKs) have historically dominated the majority of characterized natural products [30, 31], our analysis indicates that the most abundant BGC classes detected by the pipeline are terpenes and ribosomally synthesized and post-translationally modified peptides (RiPPs) (Figure 2B). This shift is likely due to recent advancements and expanded detection rules for these biosynthetic classes in state-of-the-art BGC prediction tools [14].

**Fig 2.**
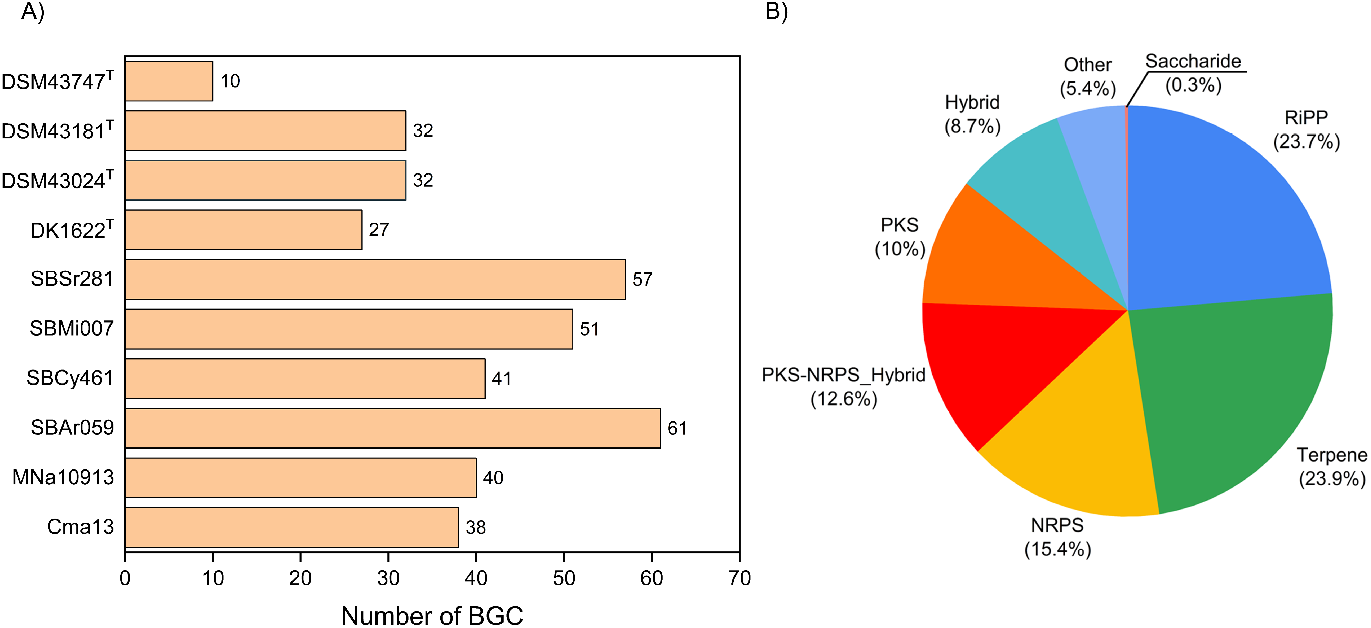
**A)** Number of BGCs predicted across each strain. **B)** Distribution of BGCs across major biosynthetic classes.

### bgc_clustering

To explore the biosynthetic diversity encoded within these genomes, the pipeline next clustered all predicted BGCs into gene cluster families (GCFs) using BiG-SCAPE [19] and BiG-SLiCE [20] tools. Reference clusters from the MIBiG database were also incorporated in the analysis to enable direct comparison with previously characterized natural product biosynthesis pathways. BiG-SLiCE identified a total of 368 GCFs across all strains, of which only 16 (4.3%) overlapped with MIBiG, while the remaining 352 (95.7%) were unique, previously uncharacterized families (Figure 3A). A similar trend was observed using BiG-SCAPE (Figure 3B), which detected 347 GCFs in total, with only 46 (13.3%) shared with MIBiG and 301 (86.7%) classified as novel.

**Fig 3.**
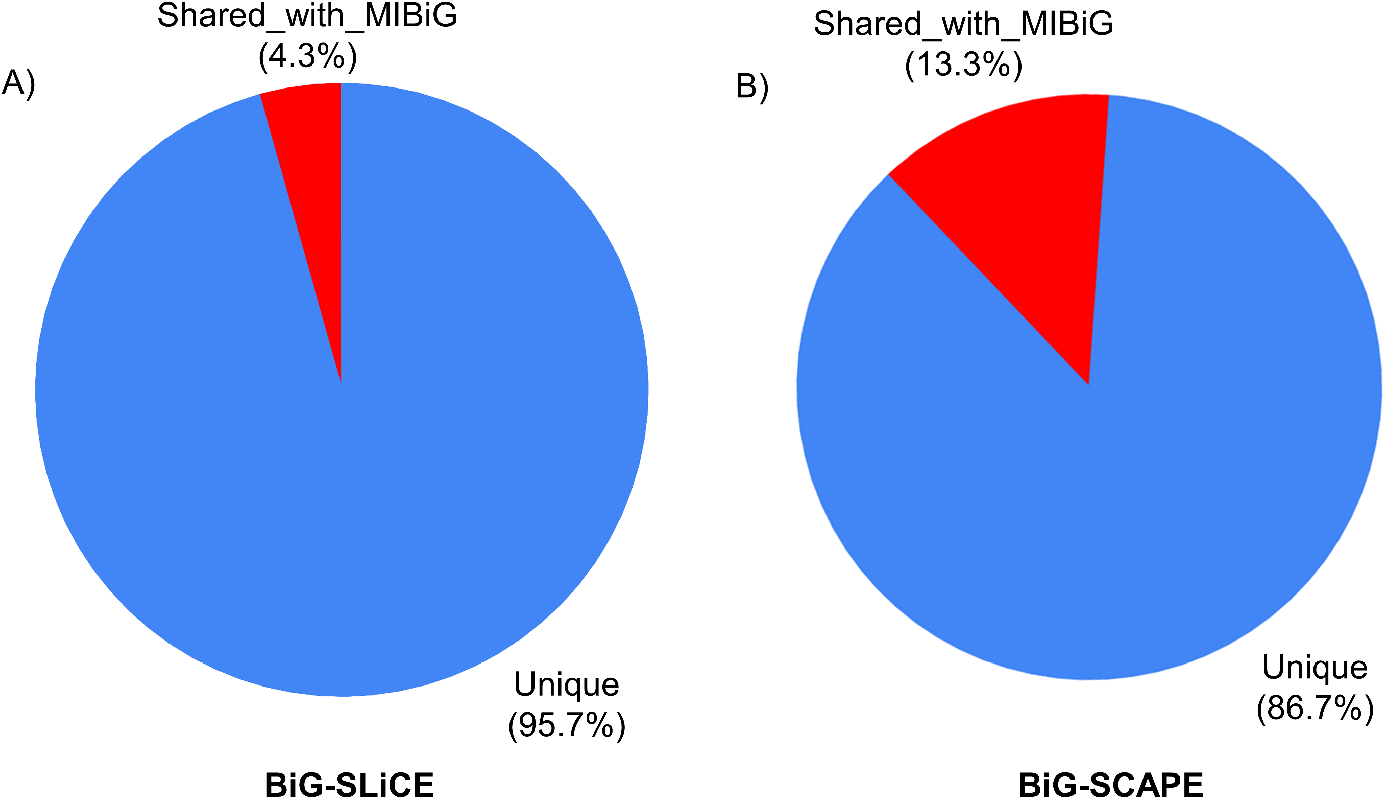
**A)** Unique and shared GCFs between the ten strains and the MIBiG database as determined by BiG-SLiCE clustering. **B)** Unique and shared GCFs between the ten strains and the MIBiG database as determined by BiG-SCAPE clustesring.

Importantly, several well-characterized MIBiG reference clusters, including myxochromide A, myxovirescin, myxoprincomide, ectoine, and others, were consistently clustered with their corresponding or related BGCs identified in our strains by both BiG-SLiCE and BiG-SCAPE. BiG-SCAPE additionally clustered the myxochelin A/B reference cluster with its corresponding BGC from DK1622^T^. Together, these results validate the ability of our pipeline to accurately group related BGCs and highlight its utility in distinguishing known pathways from novel ones, thereby facilitating efficient prioritization for natural product discovery.

## Discussion

This work introduces LoReMINE, an end-to-end, fully automated long-read based genome mining pipeline designed to streamline microbial natural product discovery. Applying the pipeline to ten bacterial strains demonstrated its ability to generate high-quality assemblies and comprehensive biosynthetic profiles. By systematically integrating multiple state-of-the-art assemblers and employing a weighted scoring framework, LoReMINE consistently produced highly contiguous circularized genomes for all strains. This level of contiguity is particularly important for detecting modular biosynthetic systems such as large PKS and NRPS clusters, where fragmented assemblies can obscure domain architecture and disrupt functional interpretation. This often leads to inaccurate GCF assignments, and hence erroneous dereplication of the resulting natural products. Clustering of predicted BGCs alongside MIBiG reference clusters revealed that well-characterized pathways were correctly grouped together, while the majority of GCFs in these strains remain uncharacterized, highlighting both the reliability of the pipeline and the vast unexplored biosynthetic diversity in these microbial genomes. These results demonstrate the value of integrated and scalable genome mining workflows for exploring and prioritizing microbial biosynthetic diversity.

A major strength of LoReMINE lies in its flexible architecture. The pipeline supports both single-strain analysis and simultaneous batch processing of multiple strains, making it suitable for small laboratory datasets as well as large comparative genomics projects. The pipeline also produces structured outputs for each submodule, which simplifies large-scale comparative analyses and supports experimental prioritization by identifying BGCs lacking similarity to known clusters from MIBiG. Furthermore, its organism-independent design enables application across diverse microbial lineages, facilitating broad exploration of microbial secondary metabolism.

Another major advantage of LoReMINE is the automation of the entire genome mining process. Traditional genome mining approaches often rely on multiple tools and lack integrated workflows, requiring extensive manual curation and expert intervention at each step, which can lead to variable results. LoReMINE streamlines this whole process by integrating all the required tools into a single automated, end-to-end workflow, minimizing manual intervention and ensuring reproducibility and reliability across large datasets.

Overall, LoReMINE provides an integrated, scalable, and reproducible framework for high-throughput microbial genome mining, enabling systematic exploration and prioritization of biosynthetic pathways to support the discovery of new natural products across diverse microbial taxa

## Conclusion

LoReMINE provides a fully automated end-to-end solution for microbial genome mining by integrating long-read genome assembly, taxonomic classification, biosynthetic gene cluster (BGC) prediction, and clustering of BGCs into gene cluster families (GCFs). By producing high-quality assemblies, the pipeline reduces the number of fragmented BGCs, enhancing confidence in further downstream analyses. Its modular design provides flexibility by allowing analyses on various scales from single strains to large datasets and making it accessible to a broad range of users. Application of LoReMINE to a diverse set of bacterial strains demonstrated its ability to uncover a wide array of largely uncharacterized BGCs, while accurately grouping known biosynthetic pathways, highlighting both the reliability of the pipeline and the considerable hidden biosynthetic diversity present in these microbial genomes. Overall, LoReMINE streamlines and standardizes genome mining workflows, enabling high-throughput, reproducible exploration of microbial secondary metabolism and supporting genomics-guided discovery of novel natural products.

## Acknowledgments

We thank Anna-Maria Kastner for sequencing the type strains used in this study. We are also grateful to Guangyi Chen for testing the pipeline and providing valuable feedback on the manuscript.

