## Supplementary information for "LoReMINE: Long Read-based Microbial genome mining pipeline"

```
===========================
| ASSEMBLY RANKING SUMMARY |
===========================
  assembly_name  has_circular_chromosome  num_circular  num_contigs       N50  total_length
0          flye                     True             1            1  12630305      12630305
1       Hifiasm                     True             1           54  12631493      16205913
2           IPA                     True             1           59  12633124      14890615

***************************************
* BEST ASSEMBLY ACCORDING TO US: flye *
***************************************

-------------------------------------------------------------------------------------
The contamination and completeness stats of the best chosen assembly is given below:
-------------------------------------------------------------------------------------
Completeness: 92.90
Contamination: 5.35
```

**Fig 1.** Example of the "`chosen_best_assembly.txt`" file generated by running the assembly using Pacbio HiFi reads. Assemblies are ranked from best to worst based on their assembly score, with the top-scoring assembly selected as the best. Completeness and contamination stats for the best chosen assembly are also included.

```
===========================
| ASSEMBLY RANKING SUMMARY |
===========================
  assembly_name  has_circular_chromosome  num_circular  num_contigs      N50  total_length
4          5000                    False             0            1  7349698       7349698
0          1000                    False             0            2  7333185       7334695
1          2000                    False             0            2  7333177       7341132
2          3000                    False             0            2  7333174       7343480
5          6000                    False             0            2  7350967       7385594
3          4000                    False             0            2  7349706       7372403
6          7000                    False             0            3  7380927       7433945
7          8000                    False             0            6  2162099       7432834
8          9000                    False             0           14  1203645       7568746
9         10000                    False             0           19   749979       7623762

***************************************
* BEST ASSEMBLY ACCORDING TO US: 5000 *
***************************************

-------------------------------------------------------------------------------------
The contamination and completeness stats of the best chosen assembly is given below:
-------------------------------------------------------------------------------------
Completeness: 98.71
Contamination: 3.23
```

**Fig 2.** Example of the "`chosen_best_assembly.txt`" file generated using the `--alt_param` option for PacBio CLR and ONT reads. Assemblies are ranked from best to worst based on their assembly score, with the top-scoring assembly selected as the best. Completeness and contamination statistics for the best chosen assembly are also included.

```
##################### NCBI database taxonomy result #####################

The identified taxonomy using NCBI is:

        1. Organism: Archangium violaceum, NCBI Accession: GCA_000733295.1, NCBI Taxonomy ID: 83451

##################### GTDB database taxonomy result #####################

The identified taxonomy using GTDB database is:

        1. d__Bacteria;p__Myxococcota;c__Myxococcia;o__Myxococcales;f__Myxococcaceae;g__Archangium;s__Archangium violaceum (NCBI accession: GCF_000733295.1)
```

**Fig 3.** Example of the "`identified_taxonomy.txt`" file for a genome where both the NCBI and GTDB databases provided a definitive species-level assignment.

```
#################### NCBI database taxonomy result ####################

Sorry, we weren't able to conclusively identify the taxonomy using NCBI. However, the top candidates (in decreasing order of priority) are listed below:

        1. Organism: Streptomyces albidoflavus, NCBI Accession: GCA_004195735.1, NCBI Taxonomy ID: 1886
        2. Organism: Streptomyces koyangensis, NCBI Accession: GCA_003428925.1, NCBI Taxonomy ID: 188770

#################### GTDB database taxonomy result ####################

The identified taxonomy using GTDB database is:

        1. d__Bacteria;p__Actinomycetota;c__Actinomycetes;o__Streptomycetales;f__Streptomycetaceae;g__Streptomyces;s__Streptomyces albidoflavus (NCBI accession: GCF_000719955.1)
```

**Fig 4.** Example of the "`identified_taxonomy.txt`" file for a genome where one database provided a definitive species-level assignment, while the other returned the closest species in decreasing order of similarity.

```
#################### NCBI database taxonomy result ####################

Sorry, we weren't able to identify the exact taxonomy using NCBI as all the hits are below the defined species threshold. However, the identified candidates (in decreasing order of priority) below the threshold are listed below:

        1. Organism: Archangium lansingense, NCBI Accession: GCA_026626635.1, NCBI Taxonomy ID: 2995310
        2. Organism: Archangium gephyra, NCBI Accession: GCA_001027285.1, NCBI Taxonomy ID: 48
        3. Organism: Archangium violaceum, NCBI Accession: GCA_000733295.1, NCBI Taxonomy ID: 83451
        4. Organism: Archangium lipolyticum, NCBI Accession: GCA_024623785.1, NCBI Taxonomy ID: 2970465

#################### GTDB database taxonomy result ####################

Sorry, we weren't able to identify the exact taxonomy using GTDB database as all the hits are below the defined species threshold. However, the identified candidates (in decreasing order of priority) below the threshold are listed below:

        1. d__Bacteria;p__Myxococcota;c__Myxococcia;o__Myxococcales;f__Myxococcaceae;g__Archangium;s__Archangium lansinium (NCBI accession: GCF_026626635.1)
        2. d__Bacteria;p__Myxococcota;c__Myxococcia;o__Myxococcales;f__Myxococcaceae;g__Archangium;s__Archangium sp035645295 (NCBI accession: GCA_035645295.1)
        3. d__Bacteria;p__Myxococcota;c__Myxococcia;o__Myxococcales;f__Myxococcaceae;g__Archangium;s__Archangium gephyra (NCBI accession: GCF_001027285.1)
        4. d__Bacteria;p__Myxococcota;c__Myxococcia;o__Myxococcales;f__Myxococcaceae;g__Archangium;s__Archangium violaceum (NCBI accession: GCF_000733295.1)
        5. d__Bacteria;p__Myxococcota;c__Myxococcia;o__Myxococcales;f__Myxococcaceae;g__Archangium;s__Archangium sp035633935 (NCBI accession: GCF_035633935.1)
        6. d__Bacteria;p__Myxococcota;c__Myxococcia;o__Myxococcales;f__Myxococcaceae;g__Archangium;s__Archangium sp036277355 (NCBI accession: GCF_036277355.1)
        7. d__Bacteria;p__Myxococcota;c__Myxococcia;o__Myxococcales;f__Myxococcaceae;g__Archangium;s__Archangium sp036388035 (NCBI accession: GCF_036388035.1)
        8. d__Bacteria;p__Myxococcota;c__Myxococcia;o__Myxococcales;f__Myxococcaceae;g__Archangium;s__Archangium violaceum_A (NCBI accession: GCF_016859125.1)
        9. d__Bacteria;p__Myxococcota;c__Myxococcia;o__Myxococcales;f__Myxococcaceae;g__Archangium;s__Archangium sp003044305 (NCBI accession: GCF_003044305.1)
        10. d__Bacteria;p__Myxococcota;c__Myxococcia;o__Myxococcales;f__Myxococcaceae;g__Archangium;s__Archangium violaceum_B (NCBI accession: GCF_016887565.1)
```

**Fig 5.** Example of the "`identified_taxonomy.txt`" file for a genome with no definitive species-level assignment from either database, but showing the closest species in decreasing order of similarity.

```
#################### NCBI database taxonomy result ####################

Sorry, we were not able to identify the taxonomy using NCBI database

#################### GTDB database taxonomy result ####################

Sorry, we were not able to identify the taxonomy using GTDB database
```

**Fig 6.** Example of the "`identified_taxonomy.txt`" file for a genome with no close matches found and no species-level assignment possible for either database.
